# Comparative and Integrative Single Cell Analysis Reveals New Insights into the Transcriptional Immaturity of Stem Cell-Derived β Cells

**DOI:** 10.1101/2023.11.15.567281

**Authors:** Mason D. Schmidt, Matthew Ishahak, Punn Augsornworawat, Jeffrey R. Millman

**Author notes:** To whom correspondence should be addressed: Jeffrey R. Millman.

## Abstract

Diabetes cell replacement therapy has the potential to be transformed by human pluripotent stem cell-derived β cells (SC-β cells). However, the precise identity of SC-β cells in relationship to primary fetal and adult β-cells remains unclear. Here, we used single-cell sequencing datasets to characterize the transcriptional identity of islets from in vitro differentiation, fetal islets, and adult islets. Our analysis revealed that SC-β cells share a core β-cell transcriptional identity with human adult and fetal β-cells, however SC-β cells possess a unique transcriptional profile characterized by the persistent expression and activation of progenitor and neural-biased gene networks. These networks are present in SC-β cells, irrespective of the derivation protocol used. Notably, fetal β-cells also exhibit this neural signature at the transcriptional level. Our findings offer insights into the transcriptional identity of SC-β cells and underscore the need for further investigation of the role of neural transcriptional networks in their development.

## Introduction

Pancreatic β-cells are the primary insulin-producing cells and therefore play a crucial role in maintaining blood glucose levels. Dysfunction or autoimmune destruction of these cells leads to diabetes mellitus, a chronic metabolic disease that is currently incurable. Directed differentiation of human pluripotent stem cells (hPSCs) into insulin-producing stem cell-derived β (SC-β) cells holds immense promise as a potentially unlimited supply of functional β-cells to treat insulin-dependent diabetes through cell replacement therapy^1–3^. This process involves a stepwise combination of small molecules, growth factors, and microenvironmental cues to drive cells through several intermediate progenitor cell types^4–7^. The resulting hPSC-derived islets (SC-islets) possess many features of primary human islets, such as a similar cell composition consisting of SC-β cells along with other islet cell types and, most notably, the ability to secrete insulin in response to glucose and restore normoglycemia in animal models. Several protocols for producing SC-islets via *in vitro* differentiation have been published^4–7^. These methods differ in many significant process parameters, including the composition of factors in the media, the types of culture vessels, and formation of the final three-dimensional aggregates. However, all of these protocols produce 3D cellular constructs that are uncontrollably heterogeneous, resulting in off-target cell populations, and are transcriptionally and functionally immature compared to their primary islet counterparts. This suggests that current *in vitro* differentiation methodologies do not fully replicate normal *in vivo* pancreatic development.

Understanding the specific pattern of gene expression that directs differentiation and maintains cell identity is critical to improving the efficiency of SC-β cell generation protocols. Recently, single-cell RNA sequencing (scRNAseq) has been applied to characterize the transcriptomic profile of SC-islets and primary human islets in various contexts^8–10^. Notably, this technology led to the identification of a substantial off-target population in SC-islets consisting of serotonin producing-cells that express genes associated with intestinal enterochromaffin cells^8^. Additionally, scRNAseq of SC-islets after transplantation into mice demonstrated that cellular identity and maturation state changes significantly *in vivo*^10–12^. While these studies have provided a comprehensive characterization of cellular identities generated by their respective protocols, no prior study has thoroughly compared cellular identities of SC-islets across different protocols. Further, the transcriptional profile of SC-islets generated by current state-of-the-art protocols has not been robustly benchmarked against the transcriptional profile of both primary adult islets and fetal islets. As a result, there is a gap in knowledge of how the transcriptional profile of SC-islets compares to that of normal human development^8,13^.

Here, we leverage published scRNAseq datasets of SC-islets from multiple protocols, both before and after transplantation, and datasets from both human adult and fetal islets to perform a novel comparative analysis of β-cell transcriptional profile across maturation states. The results provide robust definitions of the cell types produced across *in vitro* differentiation protocols and uncover commonalities and discrepancies between SC-islet development and human pancreatic development. Collectively, these data provide a resource that improves the characterization of cell identities found within SC-islets, facilitating the discovery of misexpressed genes and gene regulatory networks that can be targeted to further improve SC-β cell differentiation strategies.

## Results

### Identification of pancreatic endocrine cell types using integrated transcriptomic atlas

To understand the transcriptional maturation state of SC-islets in direct comparison to their human adult and fetal counterparts, we collected published scRNAseq datasets of human pancreatic islets from a variety of sources for comparative analysis. This included SC-islets cultured to their endpoint^8–11^, SC-islets transplanted into the kidney capsules of mice for 1 or 6 months^10,11^, primary adult islets from healthy, male and female donors age 19 to 56^14–16^, and primary fetal islets from 110 to 122 days post-conception (dpc)^17,18^ (Fig. 1a). Raw data was processed, and quality control measures were performed to remove dead cells and sequencing doublets (see Methods and Table S1). We performed unsupervised clustering on each individual dataset to generate Uniform Manifold Approximation and Projection (UMAP) plots to visualize dimensional reductions in 2D. For each dataset, clusters expressing high levels of chromogranin A (*CHGA*) were isolated as probable endocrine cell types^19,20^, narrowing down our analysis from 128,204 total pancreatic islet cells to 60,197 *CHGA+* pancreatic islet cells.

**Fig. 1.**
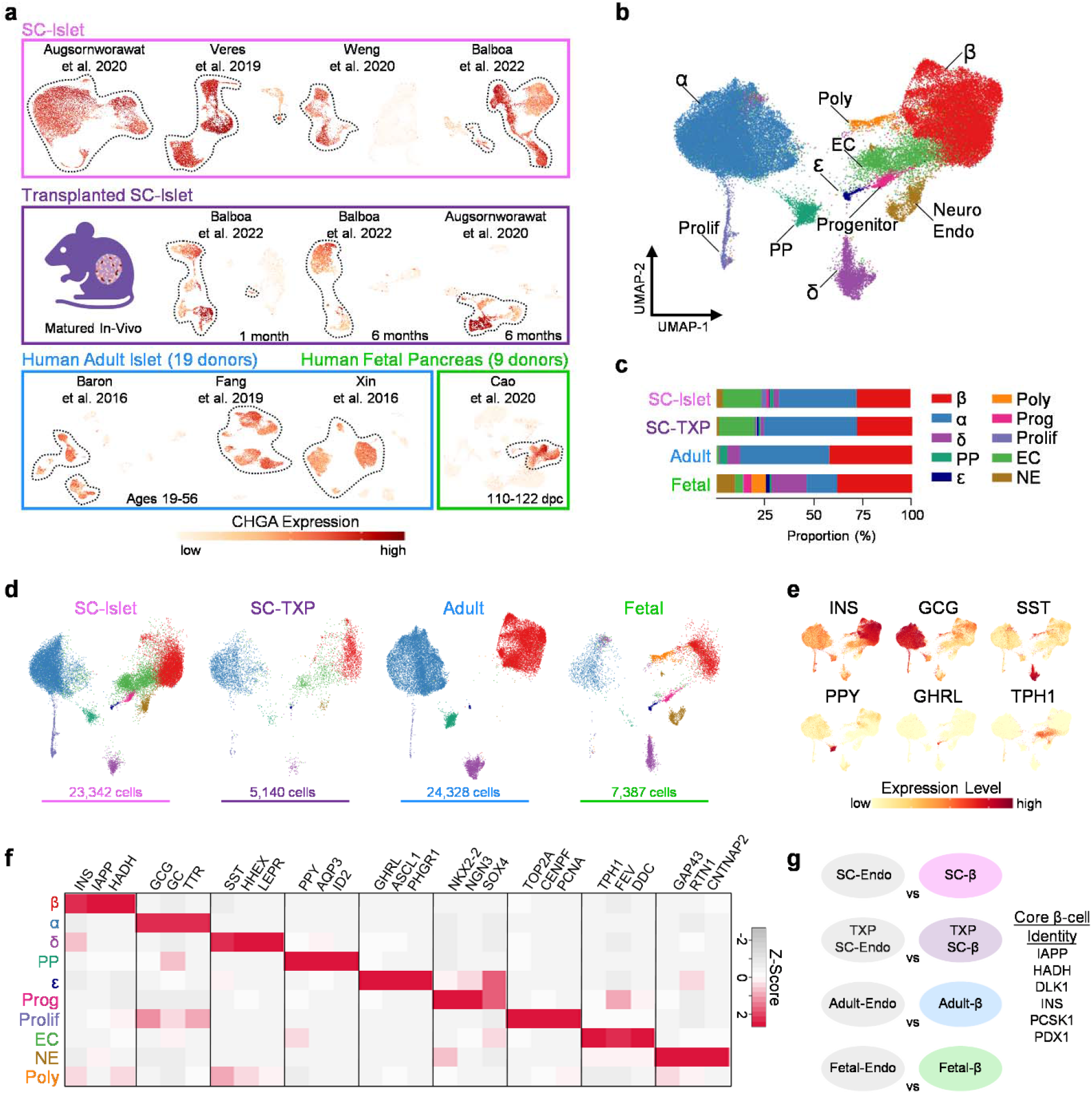
SC-islets share endocrine cell types with fetal and adult islets, see also Table S1 & Table S2. (a) Schematic of 19 human adult islets, male and female, age 19-56; 9 human fetal pancreases, male and female, 110-122 dpc; SC-islets derived from 4 unique protocols; and SC-islets derived from 2 unique protocols and transplanted into the kidney capsules of mice for 1-6 months. Each individual dataset is plotted onto a UMAP which indicates scaled expression of *CHGA* after quality control filtering. (b) UMAP of all endocrine cells integrated from each published dataset with 10 unique cell types identified. Polyhormonal (Poly), Endocrine Progenitor (Prog), Proliferating Endocrine (Prolif), Enterochromaffin-like (EC), Neuroendocrine (NE). (c) Proportion of identified cell-types from SC, SC-TXP, adult, and fetal islet sources. (d) Integrated endocrine UMAP split between SC, SC-TXP, adult, and fetal sources. (e) Feature plots indicating scaled expression level of various islet cell hormones. (f) Heatmap indicating top differentially expressed genes for each endocrine cell population. (g) Pairwise analysis indicating differentially expressed genes (log2FC > 0.3) between β-cells and all other endocrine cells shared between SC, SC-TXP, fetal, and adult islets. DEGs enriched in β-cells from all four sources make up core β-cell identity.

All *CHGA+* cells were integrated to identify shared cell populations present across each dataset. This led to the identification of 10 unique endocrine cell populations (Fig. 1b) of which the top differentially expressed genes (DEGs) are listed in Table S1. The only identifiable cell populations in the adult islets were β, α, PP, δ, ε, and proliferating endocrine, and these cell populations were also present in fetal and SC-islets (Fig. 1c-d). High expression of hormones *INS*, *GCG*, *SST*, *PPY*, and *GHRL*, along with enrichment of other cell-specific markers assisted in the identification and validation of these endocrine cell populations (Fig. 1e-f). An endocrine progenitor (Prog) cell population with enriched expression of transcription factors *NKX2-2*, *SOX4*, and *NGN3* was found to be present in both fetal and SC-islets. Consistent with previous findings^8^, a population resembling enterochromaffin-like cells (EC), marked by expression of TPH1, FEV, and DDC was only identifiable in SC-islets. Interestingly, a population of cells with neuroendocrine (NE) features, marked by enrichment of *GAP43*, *RTN1*, and *CNTNAP2*, was found to be present in both SC-islets and fetal islets. The identity and role of these endocrine cells with neuronal properties in the developing human islet has not been previously characterized. Finally, a cluster of polyhormonal (Poly) cells was identified and enriched in the fetal islets. This population is consistent with previous studies which show that cells expressing multiple hormones arise early in islet development and eventually give rise to α-cells^21,22^. This suggests the utility of this dataset to more precisely identify islet endocrine cell types than what can be surmised from the individual clustering of SC-islet scRNAseq datasets.

A universal definition of β-cell identity would not only serve as a useful resource in research, but also a potentially important attribute of cells to be used for therapy^23^. This can be particularly difficult in SC-islets, as SC-β cell identity can lack distinctiveness compared to other cell types in the tissue, particularly the SC-EC cells^8,10,24^. To establish a universal definition of healthy β-cell identity, we identified genes enriched in β-cells compared to all other endocrine cells and identified the genes whose expression is shared across all tissue sources (Fig. 1g). The β-cell genes that were most highly conserved across all sources were *INS*, *IAPP*, *DLK1*, *PDX1*, *HADH*, and *PCSK1*. We also define core identity gene lists for α, δ, and EC cells which can be found in Table S2. These gene lists provide an important definition of cell-specific islet markers that arise early in development and whose expression persists over time and across unique conditions. The strongest conserved gene signature was seen in α-cells which had a total of 32 genes that were enriched across all sources including *ARX*, *GC*, *GCG*, *IRX2*, *TTR*, and many others, while δ-cells only possessed 5 conserved identity genes (*HHEX*, *LEPR*, *SEC11C*, *SST*, and *TSHZ2*). Taken together, the assembly of an integrated pancreatic islet scRNAseq dataset with islets from human adult, fetal, and SC sources led to the precise definition of islet endocrine cell types. This dataset can serve as a tool for researchers to understand transcriptomic differences between islet cell types across unique maturation states.

### Directed differentiation protocols produce transcriptionally similar SC-islets

While several protocols for producing SC-islets have been described^25^, commonalities and differences in their transcriptional profiles are not well understood. To explore cellular heterogeneity and benchmark maturation across protocols, SC-β cells, adult-β cells, and fetal-β cells were isolated from the combined dataset and re-clustered (Fig. 2a). A detailed summary of the four differentiation protocols explored in this analysis and their associated datasets is available in Table S1. Based on clustering and Pearson correlation analysis, SC-β cells, regardless of the protocol they were derived from, appear to be transcriptionally similar when compared to adult and fetal β cells (Fig. 2b). Furthermore, SC-β cells from all protocols expressed significantly less *G6PC2*, *IAPP*, *HADH*, *UCN3*, *CHGB*, *ADCYAP1* and *SIX3* than adult-β cells (Fig. 2c). Despite their overall transcriptional similarities, unique transcriptional profiles of SC-β cells from differing protocols was still observed (Fig. 2d and Table S3). This includes SC-β cells generated by Augsornworawat, et al. expressing higher levels of *TTR*, *F10*, and *C1QL1*, while those generated by Veres, et al. express higher levels of *POTEE*, *CHGA*, and *ONECUT2*. SC-β cells generated by Weng, et al. had higher expression of *NEFM*, *AMBP*, and *NCL*, while those derived from the protocol reported by Balboa, et al. have high expression of *RPL39*, *CRYBA2*, and *CALB2*. Further work is needed to decipher if these observed differences from each protocol is important for SC-β cell function. It is important to note that Augsornworawat and Veres employed Hues8 hESC in their differentiation protocol, while Balboa and Weng employed the H1 hESC line. It is unclear whether these transcriptional differences are due to different genetic background of hESCs, culture conditions, cell preparations, and/or sequencing platforms. Despite these minute differences, the SC-β cells analyzed from these four unique datasets appear to be very similar at the transcriptional level.

**Fig. 2.**
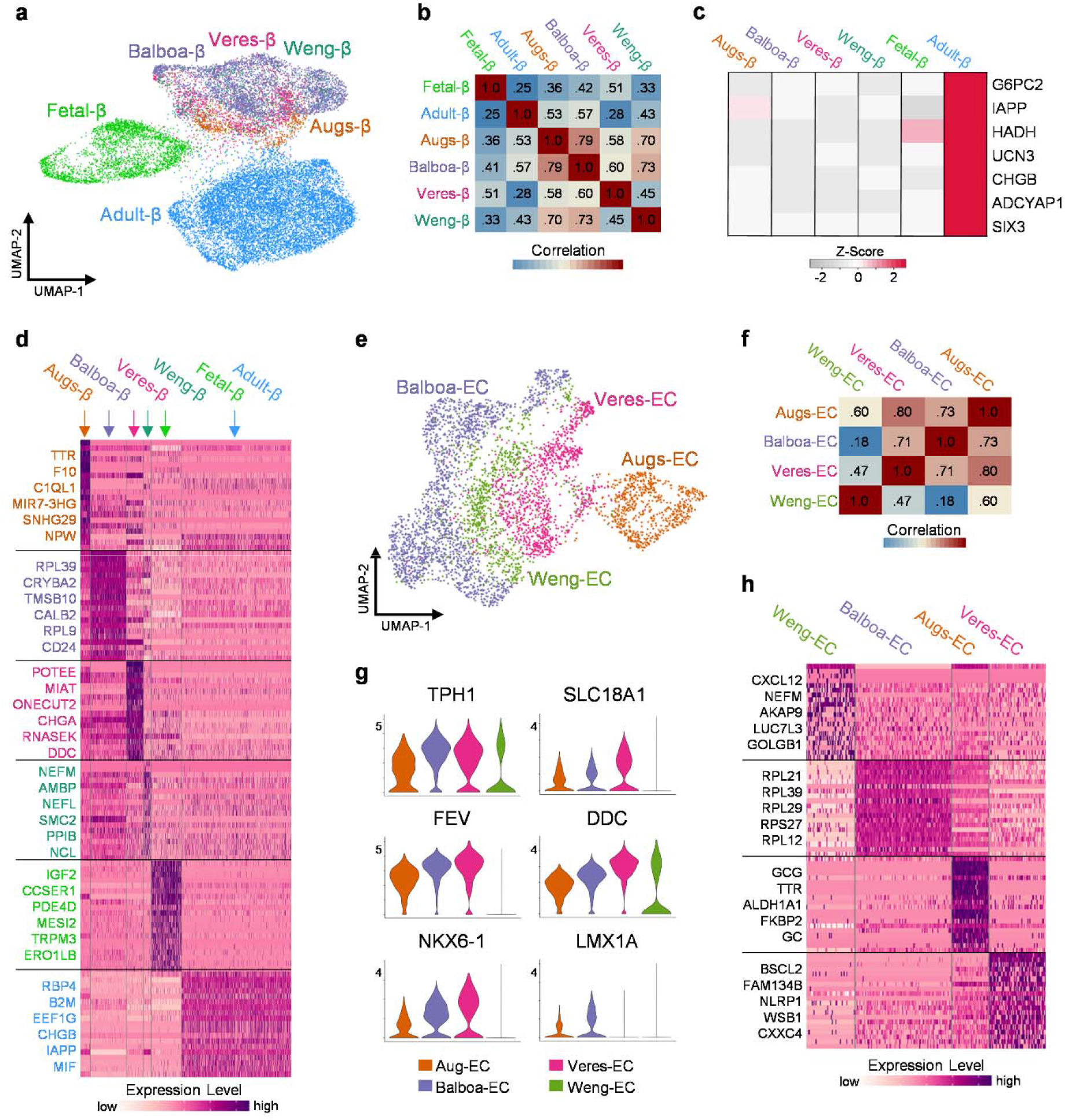
SC-β cells derived from different protocols possess similar transcriptional profiles relative to adult β cells, see also Fig. S1 & Table S3. (a) UMAP of adult β-cells, fetal β-cells, and SC β-cells clustered and split by their derivation protocol. (b) Heatmap of Pearson correlation coefficient for 1000 most variable expressed genes in β-cells. (c) Heatmap indicating average expression of β-cell maturation markers. (d) Heatmap of scaled RNA expression indicating top 20 most differentially expressed genes for β-cells derived by unique protocols, fetal β-cells, and adult β-cells. (e) UMAP of SC-EC cells clustered and split by their derivation protocol. (f) Heatmap of Pearson correlation coefficient for 1000 most variable expressed genes in SC-EC cells. (g) Violin plots indicating expression level of SC-EC identity markers across derivation protocols. (h) Heatmap of scaled RNA expression indicating top 20 most differentially expressed genes for SC-EC cells across protocols.

A similar comparative analysis was performed on SC-EC cells from each SC-islet dataset, which are marked by high expression of *TPH1*. From the combined dataset, SC-EC cells were isolated and re-clustered (Fig. 2e). The expression of key SC-EC marker genes and Pearson correlation analysis suggested that the overall transcriptional profile of SC-EC was similar across protocols (Fig. 2f-g). Analysis of DEGs revealed key differences between the SC-EC cells from different protocols (Fig. 2h). Notably, cells derived by Augsornworawat, et al. had increased expression of α-cell markers *GCG*, *TTR*, and *GC*; while SC-EC cells from Veres, et al. had the highest expression of the canonical EC-identity markers *SLC18A1*, *DDC*, and *FEV*. Interestingly, the SC-EC cells generated by Weng, et al. had the lowest expression of these SC-EC cell markers, and SC-EC cells generated by Balboa, et al. were unique for having high expression of ribosomal genes, similar to the SC-β cells from this study.

We also explored other SC-islet endocrine cell types across protocols. DEGs for SC-α, SC-δ, and SC-EC from each protocol are highlighted in Table S3. We observed few major differences in the transcriptome of SC-α and SC-δ cells from the different protocols (Fig. S1). Of note, SC-α cells from all four protocols expressed equivalent amounts of *GCG* and *TTR* to their human counterparts (Fig. S1d). Furthermore, SC-δ cells from each protocol were greatly lacking expression of *RBP4* compared to adult-δ cells (Fig. S1h). In conclusion, these results indicate that SC-endocrine cells derived from different SC-islet protocols all have similar gene expression profiles to one another, with a few notable differences. Further studies will be necessary to decipher if these transcriptional similarities in SC-islet cell types are translated to their functionality, and if the minor transcriptional differences are due in fact to differences in the differentiation protocol itself or other experimentally uncontrolled factors evident in this analysis of published datasets.

### SC-β cells are transcriptionally more mature than fetal β cells

Previous single-cell sequencing studies have shown that SC-β cells are transcriptionally immature^8–10^. We first characterized maturation in SC-β cells by comparing their global transcriptional landscape to adult and fetal β-cells (Fig. 3a). Pearson correlation of the 2000 most variably expressed β-cell genes revealed that SC-β cells had a correlation coefficient of 0.6 compared to adult β-cells and increased slightly after transplantation (Fig. 3b). Meanwhile the Pearson correlation coefficient of fetal β-cells compared to adult β-cells was just 0.33. To gain a better understanding of the unique transcriptional profile associated with SC-β cells, we performed pairwise comparisons with their primary adult and fetal counterparts. DEGs previously characterized in the context of β-cells, as well as genes with no previously identified role in β-cell identity or function were found to be enriched in either SC-β, adult-β, or fetal-β cells (Fig. 3c-d and Table S4). Interestingly, among the DEGs with log_2_(fold change) > 2 enrichment in SC-β cells were the genes *NEFM*, *CALB2*, *NEFL*, and *STMN1*, which all serve an important role in neurons.

**Fig. 3.**
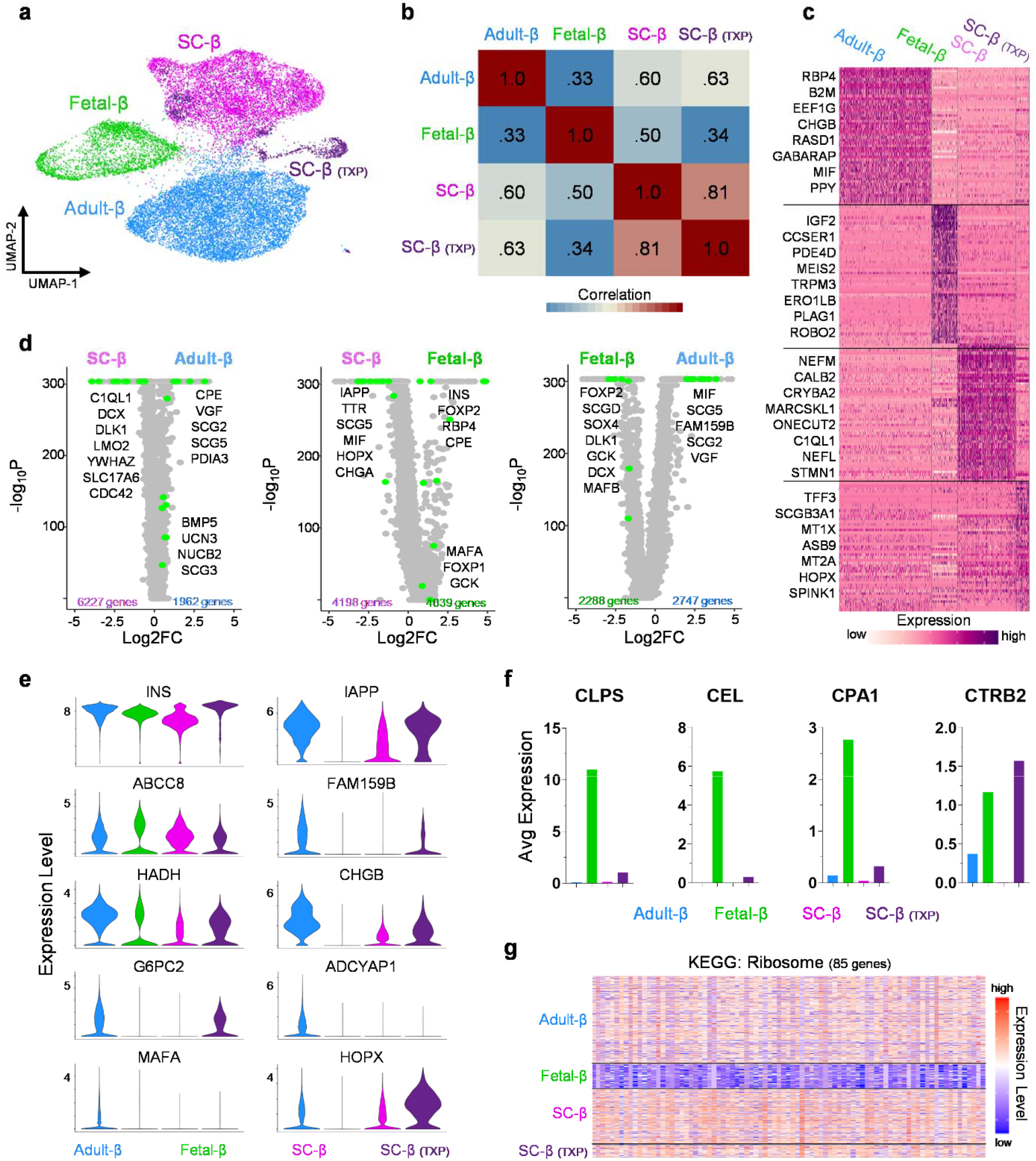
SC-β and fetal-β cells lack expression of key transcripts, see also Fig. S2 & Table S4. (a) UMAP of adult β-cells, fetal β-cells, SC β-cells, and TXP SC β-cells clustered. (b) Pearson correlation coefficient for 2000 most variable expressed genes. (c) Heatmap of scaled RNA expression indicating top 50 most differentially expressed genes for adult-β, fetal-β, SC-β cells, and TXP SC-β cells. (d) Volcano plots indicating all differentially expressed genes in SC-β vs adult-β, SC-β vs fetal-β, and fetal-β vs adult-β cells. (Adjusted p-value <0.05) (e) Violin plots indicating expression level of β-cell maturation associated genes in adult-β, fetal-β, SC-β cells, and TXP SC-β cells. (f) Bar graphs indicating average RNA counts of various exocrine markers in adult-β, fetal-β, SC-β cells, and TXP SC-β cells. (g) Heatmap indicating the scaled expression level of ribosomal genes in adult-β, fetal-β, SC-β cells, and TXP SC-β cells.

We characterized the maturation state of SC-β cells to their mature, adult counterpart by comparing the expression level of well-known β-cell maturation markers. This included *INS*, *IAPP*, *FAM159B*, *CHGB*, *G6PC2*, *ADCYAP1*, *MAFA*, and *HADH* which were all expressed at lower levels in SC-β cells compared to adult β-cells, yet most of these genes were non-existent in fetal β-cells (Fig. 3e). Transplantation of SC-islets into mice for an extended period led to the increase in expression of these maturation markers, as previously reported^10,11^. These findings show that human β-cells sourced from in-vitro differentiation of hPSCs and those sourced from primary adult and fetal islets differ in expression of a large number of genes^8,26–29^, including genes well-established to be associated with β-cell identity^30,31^. Altogether, SC-β cells lack transcriptional maturation due not only to global transcriptional disparities, but also lower expression of important β cell-maturation genes.

Our analysis also revealed that fetal β-cells possess a uniquely immature transcriptional profile. We revealed that while fetal β-cells have high expression of *INS*, they lack expression of many important β-maturation markers and have high expression of genes important for the exocrine pancreas, including *CLPS*, *CEL*, *CPA1*, and *CPA2* (Fig. 3e-f). Further, fetal β-cells have low expression ribosomal genes that are likely necessary for the production of peptides (Fig. 3g). Lastly, they have a lower fraction of cells expressing genes important for the insulin secretion mechanism (GO: 0032024) and β-cell identity^32^ compared to SC-β and adult β-cells (Fig. S2). This data supports the notion that fetal β-cells represent a transcriptional state that is less mature than SC-β cells.

### SC-β cells have persistent activity of progenitor transcription factors

Next, we set out to determine if the immature transcriptional state of SC-β cells was closely related to a β-cell progenitor state, and if we could find evidence of this progenitor state by analyzing the expression and activation of transcription factors. Therefore, we filtered our previously defined DEGs for genes that encode transcription factors and observed that both SC-β and fetal-β cells have a significantly larger enrichment of transcription factors compared to adult-β cells (Fig. 4a-b). To decipher which of these transcription factors have a role in specifying progenitor states, we filtered all expressed transcription factors for those with a previously characterized role in β-cell development. This revealed that nearly all transcription factors expressed in β cell-progenitor states are expressed in a higher percentage of SC-β cells than adult or fetal β-cells (Fig. 4c). The only exception for this was *MEIS2* which is expressed in a higher percentage of fetal and adult β-cells. To validate this observation, we generated SC-islets^33^ and compared their expression of known progenitor transcription factor to cadaveric human islets using RT-qPCR, this revealed similar trends as seen in the single-cell analysis (Fig. S3a).

**Fig. 4.**
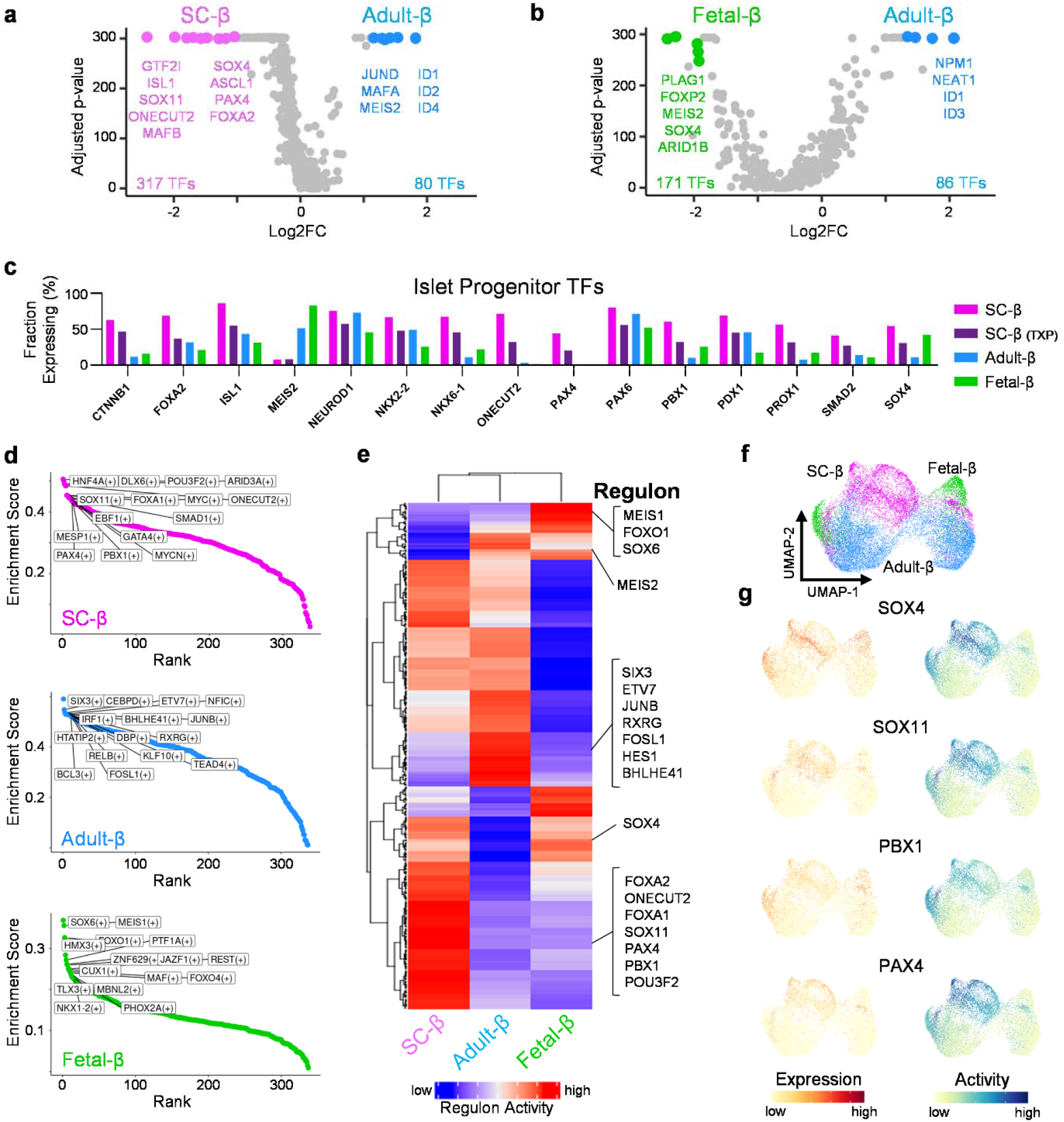
SC-β cells have high expression and activity of transcription factors associated with progenitor states, see also Fig. S3, Fig. S4, & Table S5. (a) Volcano plot indicating all expressed transcription factors between SC-β and adult-β cells (Adjusted p-value <0.05). (b) Volcano plot indicating all expressed transcription factors between fetal-β and adult-β cells (Adjusted p-value <0.05). (c) Bar plots indicating percent of cells expressing known islet developmental transcription factors in SC-β, TXP SC-β, adult-β, and fetal-β cells. (d) Chart indicating most highly enriched regulons in SC-β, adult-β, and fetal-β cells. (e) Heatmap indicating scaled regulon activity of transcription factors expressed in SC-β, adult-β, and fetal-β cells. (f) UMAP of SC-β, adult-β, and fetal-β clustered by regulon activity. (g) Feature plots indicating expression and activity of β-progenitor transcription factors.

We further explored these transcription factors by using regulon analysis^34^ to deduce and rank inferred gene regulatory networks. The most highly enriched gene regulatory networks between SC-β, adult-β, and fetal-β cells were identified (Fig. 4d and Table S5). Consistent with gene expression data, transcription factors associated with β-cell progenitor states were most highly active in SC-β cells, this includes but is not limited to: *FOXA1*, *FOXA2*, *ONECUT2*, *PAX4*, *PBX1*, *SOX4*, and *SOX11* (Fig. 4e-g). To validate the results of this regulatory gene network analysis, the expression level of the most active transcription factors and their proposed downstream targets were evaluated (Fig. S3b-c).

Next, we sought to determine if the transplantation of SC-β cells into the kidney capsule of mice for 1-month or 6-months would reduce the expression of transcription factors associated with β-progenitor states, and more closely mirror what is seen in adult-β cells. To our surprise, every transcription factor associated with β-cell development had a significantly lower expression after transplantation (Fig. S4a). To ascertain whether transcription factor activity was also reduced, we again ran transcription factor regulon analysis on SC-β and transplanted SC-β cells, and the most highly enriched regulons between both conditions were identified (Fig. S4b and Table S5). The average activity of transcription factors associated with β-progenitor states *FOXA2*, *ISL1*, *ONECUT2*, *PAX4*, *PBX1*, *PDX1*, *SOX4*, and *SOX11* all significantly decreased after transplantation (Fig. S4c). Furthermore, the decrease in expression and activity of these genes after transplantation was correlated (Fig. S4d). Collectively, these results indicate that SC-β cells have persistent expression of transcription factors associated with β-progenitor states which are reduced after transplantation. Additionally, this analysis supports that these transcription factors, especially *PAX4*, *PBX1*, *SOX4*, and *SOX11*, are still actively regulating their downstream targets.

### Dysregulated transcription factor activity drives neuronal gene program in SC-β cells

Finally, we sought to ascertain the major gene programs enriched in SC-β cells that account for their transcriptional immaturity. We employed gene set enrichment analysis (GSEA) between SC-β cells and adult β-cells and found that the top gene ontology (GO) terms enriched in SC-β cells were closely associated with neuronal morphology and function (Fig. 5a-b). β-cells have been show to share a variety of similarities with neurons including exocytotic machinery^35^, GABA containing microvesicles^36,37^, Ca^2+^ stimulated excitation^38^, neurofilament extensions^39,40^, and adhesion molecules^41^. However, the functional role these neuronal traits play in the development and function of SC-β and to the extent that they are expressed has not been previously considered.

**Fig. 5.**
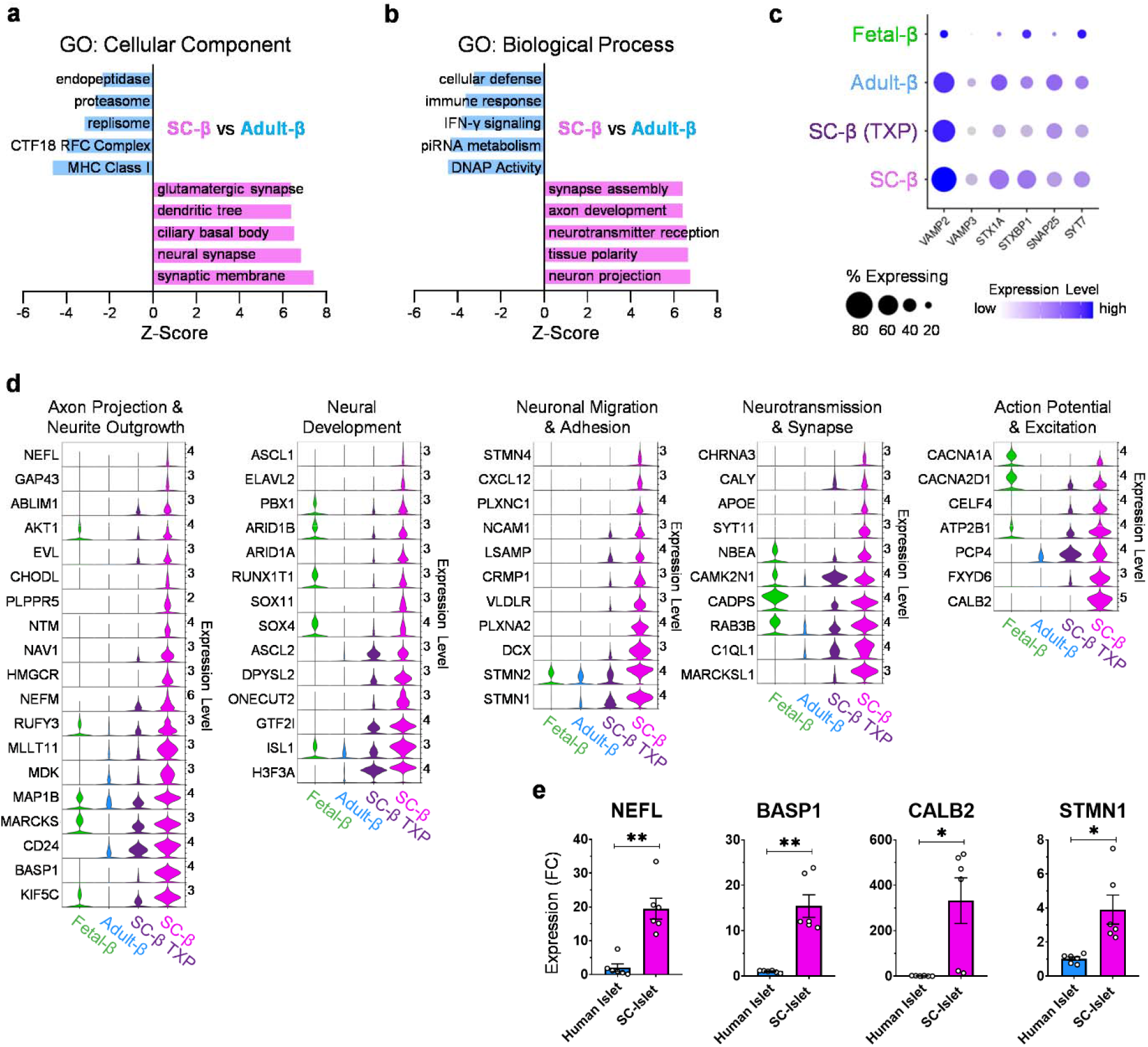
Transcripts involved in neuronal development and morphology are enriched in SC and fetal β-cells, see also Fig, S5, Fig. S6, & Table S6. (a) Bar chart indicating gene ontology: cellular component enrichment scoring of differentially expressed genes between SC-β and adult-β cells. (b) Bar chart indicating gene ontology: biological process enrichment scoring of differentially expressed genes between SC-β and adult-β cells. (c) Dotplot indicating expression level of genes associated with insulin granule exocytosis. (d) Panel of violin plots indicating expression level of genes (>1 log2FC of SC vs adult β) associated with various neuronal traits between SC, SC-TXP, adult, and fetal β-cells. (e) RT-qPCR of SC-islets (n=6) at s6d7 and human cadaveric islets (n=6) indicating fold change (FC) expression relative to TBP. All data are represented as the mean, and all error bars represent the s.e.m. Individual data points are shown for all bar graphs. ns, not significant; *P<□0.05, **P<□0.01.

Exocytosis of insulin-containing granules in β-cells is essential to their function, and many of the genes involved in this process have a similar role in neurotransmission. Therefore, we investigated the expression of *VAMP2*, *VAMP3*, *STX1A*, *STXBP1*, *SNAP25*, and *SYT7*, which are the essential components of the insulin-exocytosis machinery, yet we saw no differences in their expression between SC and adult β-cells (Fig. 5c). To further investigate the neuronal traits enriched in SC-β cells, we analyzed the expression level of large gene sets associated with axonal, synaptic, and dendritic morphology and observed their broad overexpression in SC-β cells compared to adult β-cells (Fig. S5a). Our analysis also revealed that, compared to both adult and fetal β-cells, SC-β cells overexpress genes encoding for neurofilaments and their associated proteins involved in axon guidance, genes needed in neural migration, genes essential for neurotransmission, and genes necessary for generating and maintaining action potentials (Fig. 5d). Of interest, SC-β had significant enrichment in the expression of genes that play a major role in neuronal development including *GTF2I*, *ASCL1*, *SOX4*, and *SOX11*. These genes were compiled into a curated list which defines the neuronal program that is overly enriched in SC-β cells (Table S6). We validated this observation by generating SC-islets^33^ and compared their expression of neuronal markers to cadaveric human islets using RT-qPCR, confirming that SC-islets expressed neuronal genes at a significantly higher level than adult islets (Fig 5e).

To decipher if these neuronal traits expressed widely in SC-β cells are biologically relevant or simply the effect of their *in-vitro* differentiation environment, we examined the neuronal gene program present in fetal β-cells. Similarly, when compared to adult β-cells, fetal β-cells are enriched for GO terms associated with neuronal morphology and function (Fig. S5b-c). Furthermore, they also contain higher expression of genes necessary for the formation of synapses, axons, and dendrites when compared to adult β-cells. We also found that SC-β cells that had been transplanted into mice showed loss of these previously described neuronal properties. To further validate that this neural gene program was not a result of cell-lines used, differentiation protocol, or sequencing platform we analyzed three additional datasets. These additional analyses showed that SC-β derived with induced pluripotent stem cells (iPSC)^10^ and other cell-lines^42^, as well as the use of the single-nuclei sequencing method^12^ all shared the same neural gene program when compared to human adult β-cells (Fig. S6). Furthermore, we discovered that this neural gene program is also active in SC-EC cells (Fig. S5d-f). Despite the fact that neonatal and adolescent β-cells produce serotonin^43,44^, we confirmed that SC-β cells do not express any genes associated with serotonin production. Therefore, we concluded that the population of SC-β used throughout the analysis were not contaminated with enterochromaffin-like cells. Yet, the fact that this neural gene program is shared between SC-β and SC-EC cells is an important finding and suggests that this dysfunctional neuronal gene program in SC-β cell development may be relevant to the generation of EC cells during directed differentiation. All of this suggests that a neuronal gene program is a biologically relevant phenomenon of immature β-cells, and its removal is essential for the maturation of the SC-β cell transcriptional landscape.

Several transcription factors and gene regulatory networks are shared in both β-cells and neurons during development^45^. We found that the transcription factors shared in both pancreas and neuron development are more highly expressed in SC-β cells than fetal or adult β-cells, especially genes of interest *PBX1*, *SOX4*, and *SOX11* (Fig. 6a). To see if persistent activity of progenitor associated transcription factors are activating neural gene programs in SC-β cells, we systematically analyzed the target genes of those transcription factors that are conserved in both pancreas and neuron development looking to see if they were enriched in SC-β cells. We found that the transcription factors *PBX1*, *SOX4*, *PAX4*, *ISL1*, *SOX11*, *SMAD1*, *NKX2-2*, and *DNMT3A* all possessed gene targets involved in neuronal gene programs which were highly active in SC-β cells and not present in adult β-cells (Fig. 6b). Furthermore, for *PBX1*, *SOX4*, *SOX11*, and other transcription factors, we isolated their top 50 most expressed targets in SC-β cells. When these 50 genes were analyzed with EnrichR^46–48^, the most common GO terms include axonal growth cone, synaptic vesicle membrane, neurofibrillary tangle, synaptic membrane, dopamine secretion, dendritic transport, and other cellular and biological processes in neurons (Fig. S7). Lastly, the previously curated list of genes defining the neuronal program enriched in SC-β was cross referenced with the target genes of all active transcription factors in β-cells to determine likely transcription factors that are involved in activating this neural program. The transcription factors with the most predicted targets were nearly all enriched in SC-β cells and previously implicated *PAX4*, *SOX11*, *SOX4*, and *PBX1* shared some of the most target genes with our newly defined SC-β neuronal gene program (Fig. 6c). We are not surprised to find that β-cells express neural transcription factors that play a role in pancreas development, however we find it interesting that these transcription factors are highly enriched in SC-β cells and are likely contributing to gene regulatory networks that drive a neuronal transcriptional program in SC-β cells.

**Fig. 6.**
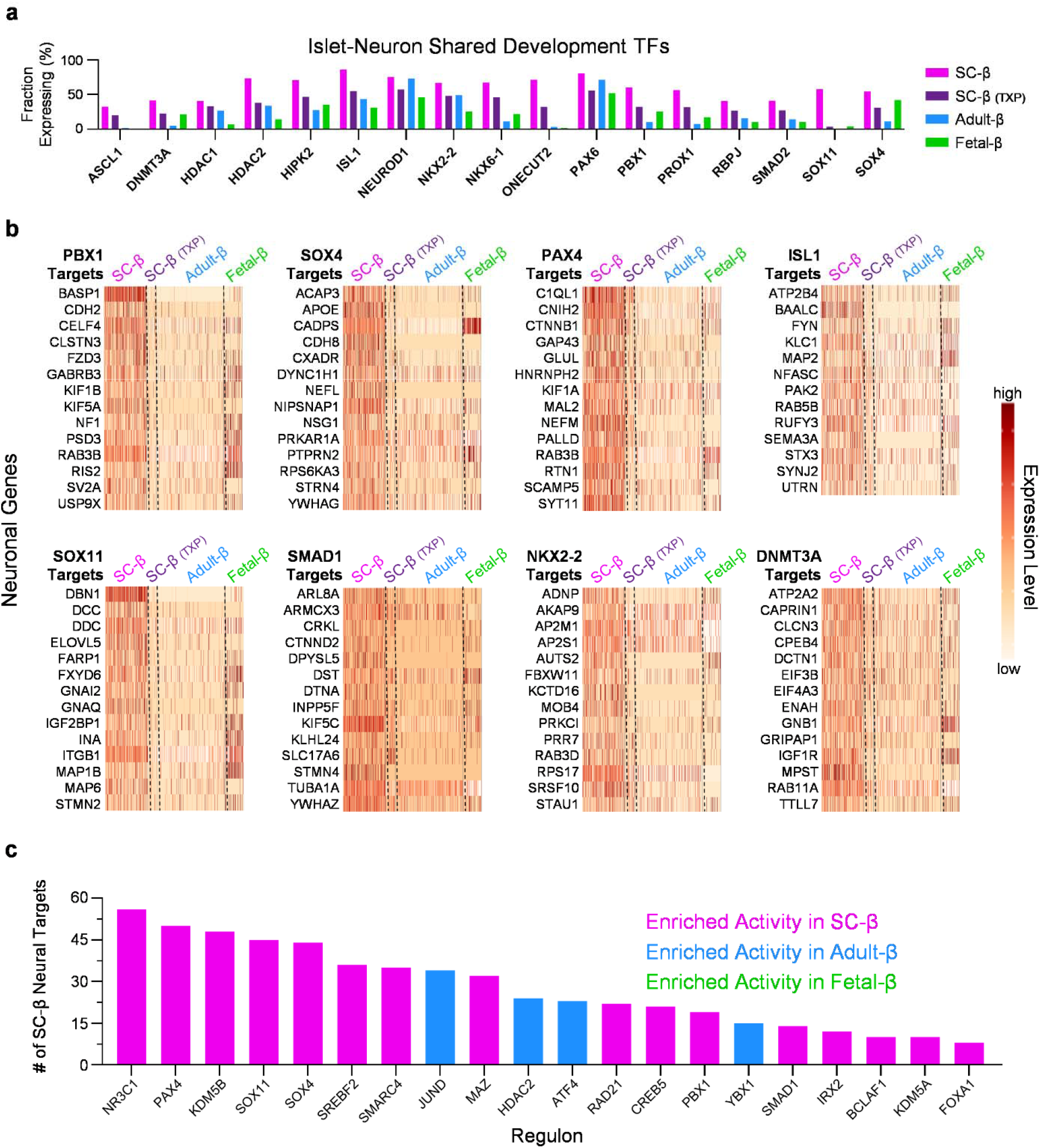
Transcription factors involved in progenitor β-cell states drive neuronal gene programs, see also Fig. S6 & Table S6. (a) Bar plots indicating percent of cells expressing transcription factors that have a role in both islet and neuron development in SC-β, TXP SC-β, adult-β, and fetal-β cells. (b) Panel of heatmaps indicating expression level of genes that are associated with neuronal traits and are targets of SC-β enriched regulons. (c) Bar plot indicates transcription factors active in β-cells with the most targets of SC-β neural genes (65 genes in total).

## Discussion

As SC-islets have the potential to functionally cure diabetes and move into clinic trials, the importance of understanding what defines islet and β-cell identity and maturation has become increasingly important^25^. Improving our understanding of the differences and commonalities in lineage specification could enhance the differentiation of hPSCs into islet cells, thereby boosting the efficacy of SC-islet therapy. The present study aimed to integrate multiple publicly available datasets to identify and characterize islet endocrine cell types, including β, α, and δ cells from SC-islets, fetal islets, and adult islets. This integrated dataset provides a detailed characterization of β-cell and all other islet cellular identities across a wide range of contexts. This not only distinguishes each cellular identity but also provides this information across tissue sources. The genetic programs described here could be targeted to better understand the acquisition of β-cell identity and to improve differentiation and maturation of SC-β cells during *in vitro* differentiation.

This analysis demonstrated that SC-β cells were transcriptionally more similar to adult rather than fetal β-cells. This finding is consistent with prior microarray and flow cytometry based-analysis^26,29^ but provides much greater detail and rigor for β-cell and other cell types. Although fetal β-cells express *INS*, they lack other important β-cell machinery and ribosomal genes, as the cells used here are only from around 100 dpc. Prior work has identified that fetal β-cells have higher expression of *ISL1*, *NEUROG3*, and genes associated with immune function compared to adult β-cells^49^, and fetal islet maturation is marked by loss of polyhormonal state and decreased proliferative capacity, which occurs at approximately 15 weeks post-conception (wpc)^50^. Like SC-β cells^11,33^, fetal β-cells are functionally immature compared to adult β-cells^26^, consistent with the immature gene expression signature observed in this study. Despite the heterogeneity observed in β-cells, we were able to discover that β-cells possess a core β-cell identity when compared to other endocrine cell types, consisting of expression of *INS*, *IAPP*, *DLK1*, *PDX1*, *HADH*, and *PCSK1*. We believe the β-cell and other identities defined here will be helpful for cellular identification in the field and complements prior efforts that have provided definitions of β-cells in primary tissues^32,51^.

Several groups have reported differentiation protocols that produce SC-islets^25^. This study is focused on publicly available scRNAseq datasets^8–11^ based on protocols first published by the Rezania/Kieffer^28^, Melton^29^, Millman^52^, and Otonkoski^11^ groups. While there are considerable differences in the reported *in vitro* and *in vivo* function of cells produced by these protocols, the comparability of transcriptional identities of the final cellular populations was unclear in the literature. Our analysis reveals that in general the transcriptional identities of all major cell types, including SC-β cells, was largely similar among all the protocols. This includes very low expression of *MAFA* and *UCN3*, indicating that development of protocols that can generate cells expressing high levels of these maturation markers *in vitro* is still lacking in the field. All differentiation protocols also produced enterochromaffin-like cells that were absent in fetal and adult primary tissue samples. Furthermore, there may indeed still be differences in the chromatin accessibility signature of cells produced from different *in vitro* differentiation protocols, which we expect to be answered in the near future as this line of investigation is gaining in attention^12,42^.

This study also found that that SC-β cells differed from adult β-cells through expression of neuronal and progenitor transcriptional programs. While a substantial fraction of genes are normally expressed by both β-cells and neurons^53^, such as synaptic-like microvesicles^36^ and gamma aminobutyric acid (GABA)^54^, the extent of expression of these genes is greatly elevated in SC-β and fetal β-cells. Furthermore, SC-β cells had enriched expression of progenitor-associated transcription factors^55–57^, such as *PAX4*, *PBX1*, *SOX4*, and *SOX11*, and these transcription factors were predicted to be among the most active in SC-β cells. Interestingly, SOX4 and SOX11 are also of critical importance in pan-neuronal protein expression^58^. Future studies could look at the relationship of these transcriptional identities to epigenetic states, as recent papers have demonstrated the importance of chromatin accessibility on SC-β cell identity^12,42^ and another prior study has shown that that pancreatic β-cells exhibit an active chromatin signature similar to neural tissues that appear to be dynamically regulated by Polycomb repression programs^59^.

This resource will serve as a tool for hypothesis generation in hopes of further optimizing protocols for the generation of SC-β cells. Future studies should work to understand the effects of perpetual expression and activation of progenitor transcription factors on SC-β cell function and if it is possible to enhance the maturation of SC-β cells by inhibition of these progenitor transcriptional networks. Furthermore, persistent activity of progenitor transcription factors in SC-β cells should be investigated to determine if they are responsible for the abnormal neural gene network identified in this study. To this point, further studies are needed to understand to what extent this neural gene program, ever present in SC-β cells, is translated to their functional properties. In addition, our finding that SC-EC and SC-β cells, despite being distinct cell types, share commonalities in this irregular neural gene program is interesting and presents the hypothesis that this dysregulated neural transcriptional profile present in SC-β development may contribute to the generation of SC-EC cells. Lastly, while transplantation of SC-β cells greatly refines the transcriptional profile of these cells, the mechanisms by which this is achieved still needs to be worked out.

Our analysis provides novel insights into the identity and characteristics of islet endocrine cells and highlights the importance of SC-β cells in understanding development and function. The findings contribute to a better understanding of the differences and similarities between SC, fetal, and adult islet cells and shed light on the potential of SC-β cells in diabetes treatment. We hope that these findings will allow for future studies using more robust hypothesis impacted by our novel findings.

### Limitations of Study

A limitation of this study is that we relied on published datasets for our analysis. This was done because we believed that a comprehensive and rigorous analysis and comparison of the best-in-class single-cell RNA sequencing data would lead to novel insights into islet identity and transcriptional regulation.

### Resource Availability

#### Lead Contact

Further information and requests for resources and reagents should be directed to and will be fulfilled by the lead contact, Jeffrey R. Millman.

#### Material Availability

This study did not generate new unique reagents.

#### Data and Code Availability

This paper analyzes existing, publicly available data. These accession numbers for the datasets are listed in Table S1. The Seurat object containing the integrated CHGA+ cell populations from all datasets, which is required to reproduce figures, is deposited at the Washington University Research Data (WURD) repository in standard RDS format. All other data supporting the findings of this study are available from the corresponding author on reasonable request. Codes used for integrating and analyzing scRNAseq datasets are available on https://github.com/mschmidt22. Any additional information required to reanalyze the data reported in this paper is available from the lead contact upon request.

### Experimental Procedures

#### scRNAseq Datasets

Healthy pancreatic islet scRNAseq datasets from primary adult, primary fetal, stem-cell derived islets, and transplanted stem-cell derived islets, were compiled from multiple published sources^8–11,14–18^. Primary adult islet datasets from 19 donors, 15 male and 4 female aged between 19 and 56 years of age were obtained from GSE84133, GSE101207, and GSE114297. Primary fetal islets datasets from 9 donors, 3 male and 6 female aged between 110 and 122 days post conception were obtained from of the Human Gene Expression Development Atlas (dbGaP accession number phs002003), generated and analyzed by the laboratories of Drs. Ian Glass, Jay Shendure, and Cole Trapnell, supported by funding from the National Institutes of Health to Dr. Glass (HD000836), Brotman Baty Institute for Precision Medicine to Dr. Shendure and Dr. Trapnell, the Paul G. Allen Frontiers Foundation to Dr. Shendure and Dr. Trapnell, and the Howard Hughes Medical Institute to Dr. Shendure. SC-islet datasets, in-vitro and those transplanted, were obtained from GSE151117, GSE114412, GSE143783, and GSE167880. All SC-islet datasets employed were accumulated from SC-islets that had been cultured to their mature endpoint. All other information pertaining to the raw data employed in this analysis can be found in Table S1.

#### Quality Control of Single-cell Datasets

RStudio [v1.3.1093] running R [v4.0.3] and the Seurat [v4.3.0]^60^ package were used to perform all initial analyses. Imported datasets were aligned and annotated with the reference human genome (hg38) from the EnsDb.Hsapeins.v86 database^61^. Poor quality cells including dead cells, doublets and poorly sequenced cells were excluded from this study. Briefly, dead, or apoptotic cells were excluded by filtering out cells containing high mitochondrial counts. Doublets were excluded by removing cells with exceedingly high RNA counts. Poorly sequenced cells were removed by excluding cells with low unique RNA features and low total RNA features. Thresholds for filtering poor quality cells of each individual dataset can be found in Table S1. Datasets obtained from SC-islet cells transplanted into mice required an additional removal of host cells via exclusion of cells expressing TTC36, a kidney gene that aligns to both the mouse and human genome. When applicable, meta data information including original dataset, donor age, donor BMI, and donor gender were added.

#### Integration of Datasets and Identification of Endocrine Cell Types

Subsequently, we performed integration and normalization using the Seurat [v4.3.0]^60^ package. Gene expression data was processed using *ScaleData* and *NormalizeData* to adjust gene counts. Each scRNAseq dataset was individually clustered employing the standard workflow. Briefly, clustering was performed using *FindNeighbors* and *FindClusters* with 20 dimensions and resolutions ranging from 0.4 - 4.5 to determine clusters. Cell types were identified by performing differential gene expression analysis using *FindAllMarkers.* Clusters of cells with high expression of endocrine marker gene *CHGA* were isolated using *subset* for further analysis. Fetal islets contained a population of acinar cells with expression of *CHGA*, these cells were not included in further analysis.

Some datasets did not contain mitochondrial genes, therefore mitochondrial genes were removed from all datasets, prior to integration and downstream analysis. Integration of endocrine cells from each dataset was performed by combining subset endocrine datasets into a single Seurat object using *FindIntegrationAnchors.* Cell types from multiple datasets were assigned based on the 2000 most variably expressed genes. Clustering was performed using *RunPCA* and *FindClusters* with parameters adjusted to a resolution of 2 and dimensions of 30. The top genes that separate each cluster within the integrated islet UMAP were identified with *FindMarkers* and these gene lists, included in Table S1, were used to designate the different islet endocrine cell types. Endocrine cell type identifiers were added to metadata.

#### Comparative Expression Analysis

Differential gene expression analyses comparing cell types of various conditions were computed using the wilcox test method of *FindMarkers*. The expression level of differentially expressed genes were visualized using *FeaturePlot, DoHeatmap, VlnPlot,* and *DotPlot.* Volcano plots were generated by performing differential gene expression analysis across two conditions and using *EnhancedVolcano* of the EnhancedVolcano [v1.8.0]^62^ package. Heatmaps indicating average expression were generated by computing the average values across two or more conditions using *AverageExpression* and visualized with the *heatmap.2* function of gplots [v3.1.3] package.

#### Inferred Gene Regulatory Network Analysis

To perform inferred regulatory gene network analysis, we employed the SCENIC [v0.9.18] command line interface (CLI) to construct gene regulatory networks from our scRNAseq data^34^. A loom object was created from the Seurat object which includes raw RNA counts and the assigned metadata of each cell. This loom object was used as input for the CLI workflow to score network activity. Candidate regulons, which includes a list of transcription factors for hg38 along with motif annotations and rankings, were downloaded from cisTargetDB (https://resources.aertslab.org/cistarget/). The activity of each regulon was calculated using area under the curve (AUC) calculations to assess significant recovery of a set of genes for individual cells. To generate regulons enriched in one group of cells a regulon specificity score (RSS) was computed. RSS and scaled expression of regulon activity was visualized in R using the *plotRSS_oneSet* and *ComplexHeatmap* functions.

#### Gene Set Enrichment Analysis

Gene set enrichment analyses were performed using the singleseqgset [v0.1.2.9000] package (https://github.com/arc85/singleseqgset) and the EnrichR interactive website^46^. For singleseqgset package, we used variance inflated Wilcoxon rank sum testing to determine enrichment of gene sets across specified conditions. All ontology gene sets in the Human MSigDB Collection^63–65^ were tested. For analysis using EnrichR, combined enrichment scores were computed and visualized based on Gene ontology gene sets. Combined enrichment scores were computed using Fisher exact test and multiplying that by the z-score of the deviation from the expected rank.

#### SC-islet Differentiation

The HUES8 (RRID: CVCL_B207) human embryonic stem cell (hESC) line (authenticated August 2022) was provided by Douglas Melton (Harvard University)^29^. All hESC work was approved by the Washington University Embryonic Stem Cell Research Oversight Committee (approval no. 15-002) with appropriate conditions and consent. Hues8 cells (passage 78) were removed from liquid nitrogen, unthawed, and plated with mTeSR1 (StemCell Technologies; 05850) which was used for the culture of undifferentiated stem cells. All cell culture was maintained in a humidified incubator at 5% CO2 and 37□°C. Cells were passaged every 4□days by washing cell with phosphate-buffered saline (PBS) and incubating with TrypLE at 0.2□ml□cm^−2^ (Gibco; 12-604-013) for 10□min or less at 37□°C. Dispersed cells were then mixed with an equal volume of mTeSR1 supplemented with 10□µM Y-27632 (Pepro Tech; 129382310MG). Cells were counted on Vi-Cell XR (Beckman Coulter) and spun at 300g for 3□min at room temperature (RT). The supernatant was aspirated, and cells were seeded at a density of 0.8□×□10^5^ cm^−2^ for propagation onto Matrigel (Corning; 356230)-coated plates in mTeSR1 supplemented with 10□µM Y-27632. After 24□h, medium was replaced daily with mTeSR1 without Y-27632. SC-islet differentiation was performed as described previously^33^. Briefly, hESCs were seeded at a density of 6.3□×□10^5^ cells□cm^−2^. Twenty-four hours later, the mTeSR1 was replaced with differentiation medium supplemented with small molecules and growth factors.

#### SC-islet and primary islet culture

After 7□days in stage 6 of the differentiation protocol, cells were dispersed from the culture plate with TrypLE (Gibco; 12-604-013) for up to 10□min at 37□°C. The cells were mixed with an equal volume of stage 6 enriched serum-free medium (ESFM), centrifuged at 300g, and resuspended in ESFM at a concentration of 1 million cells□ml^−1^. Five milliliters of this solution were pipetted in each well of a six-well plate and placed on an orbital shaker (Orbi-Shaker CO2, Benchmark Scientific) at 115□r.p.m. to form SC-islet clusters. These clusters were maintained by aspirating and replacing 4□ml of ESFM every 2□days. Primary human islets were acquired as clusters and shipped from Prodo Laboratories, which required consent from the donor’s relatives for use in research. Consent information can be found on their website (https://prodolabs.com/ human-islets-for-research). These islets have been refused for human islet transplants and meet specific criteria for research use. Our study consists of six donors. Upon arrival, islets were transferred into a six-well plate on an orbital shaker at 115□r.p.m. and maintained with 4□ml per well of CMRL1066 Supplemented medium (Corning; 99-603-CV) with 10% heat-inactivated fetal bovine serum (Gibco; 26140-079).

#### Real-Time qPCR

RNA was extracted from primary islets 2 days after arrival and from SC-islets (Hues8 passage 80) at s6d7 with the RNeasy Mini Kit (74016, Qiagen). Samples were treated with a DNase kit (79254, Qiagen) during extraction. The High Capacity cDNA Reverse Transcriptase Kit (4368814, Applied Biosystems) was used to synthesize cDNA on a thermocycler (A37028, Applied Biosystems). The PowerUp SYBR Green Master Mix (A25741, Applied Biosystems) was used on a QuantStudio™ 6 Pro Real-Time PCR System (A43180, Applied Biosystems), and real-time qPCR results were analyzed using a ΔΔCt methodology. TBP was used as a housekeeping gene. Primer sequences were as follows:

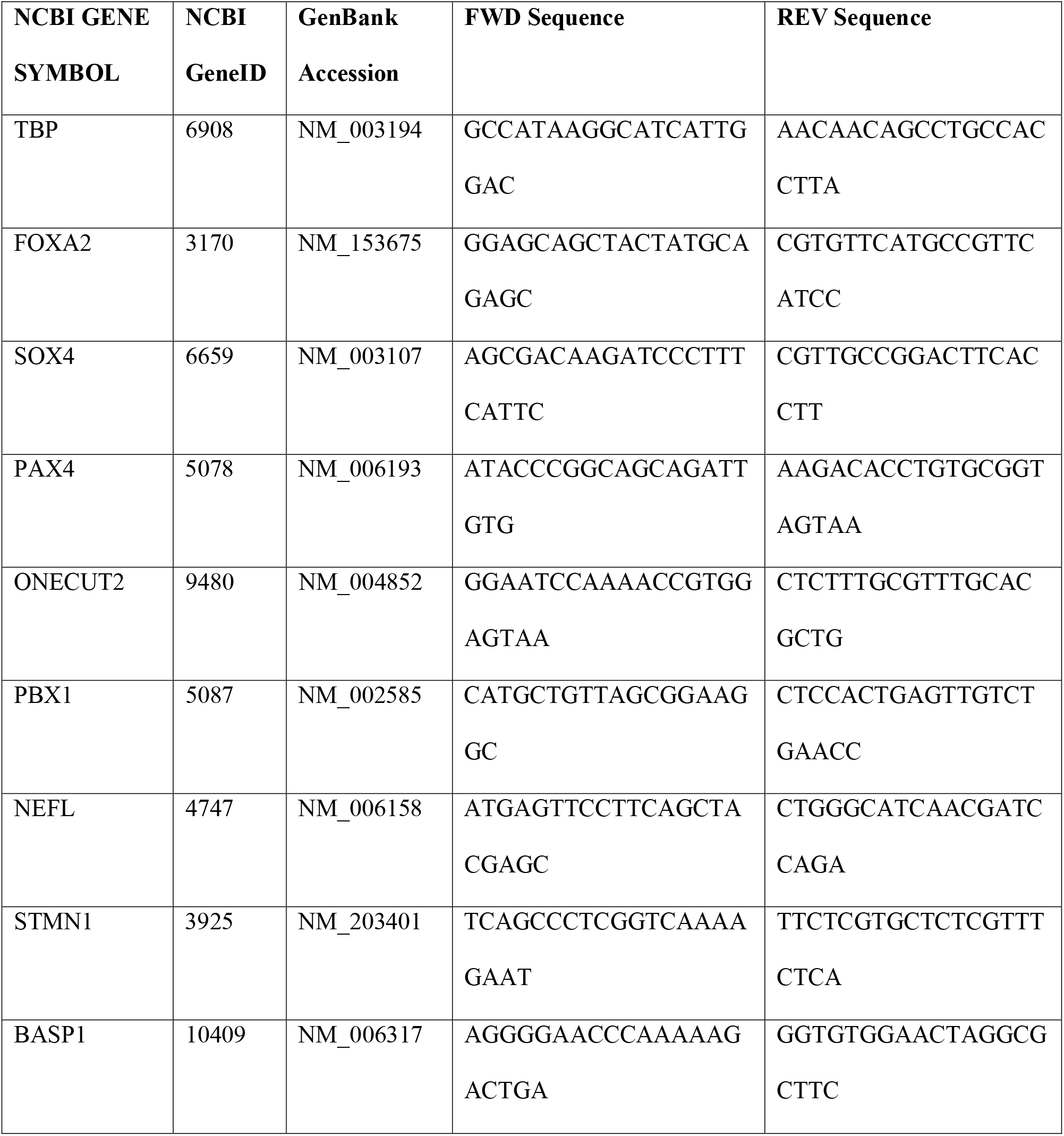
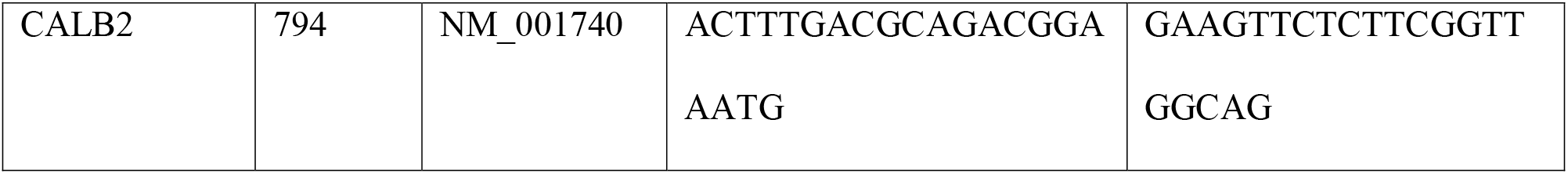

## Statistics

Statistical analysis was performed by 2-tailed unpaired t test calculated by GraphPad Prism (8.0.1). All data are presented mean ± SEM. p < 0.05 was considered statistically significant. Data analysis was not blinded.

## Acknowledgements

This work was funded by the NIH (R01DK127497), JDRF (3-SRA-2023-1295-S-B), a Human Islet Research Network (HIRN) Catalyst Award, the Edward J. Mallinckrodt Foundation, and startup funds from the Washington University School of Medicine Department of Medicine. M.I. was supported by Rita Levi-Montalcini Postdoctoral Fellowship in Regenerative Medicine and the NIH (T32DK007120).

## Author Contributions

M.D.S. and J.R.M. conceived of all computational analysis. M.D.S. performed all computational analysis. M.I. and P.A. provided vital information on all aspects of the project. M.D.S., M.I., and J.R.M. wrote the manuscript. All authors revised and approved the manuscript.

