## Supplemental Figures for "Comparative and Integrative Single Cell Analysis Reveals New Insights into the Transcriptional Immaturity of Stem Cell-Derived β Cells"

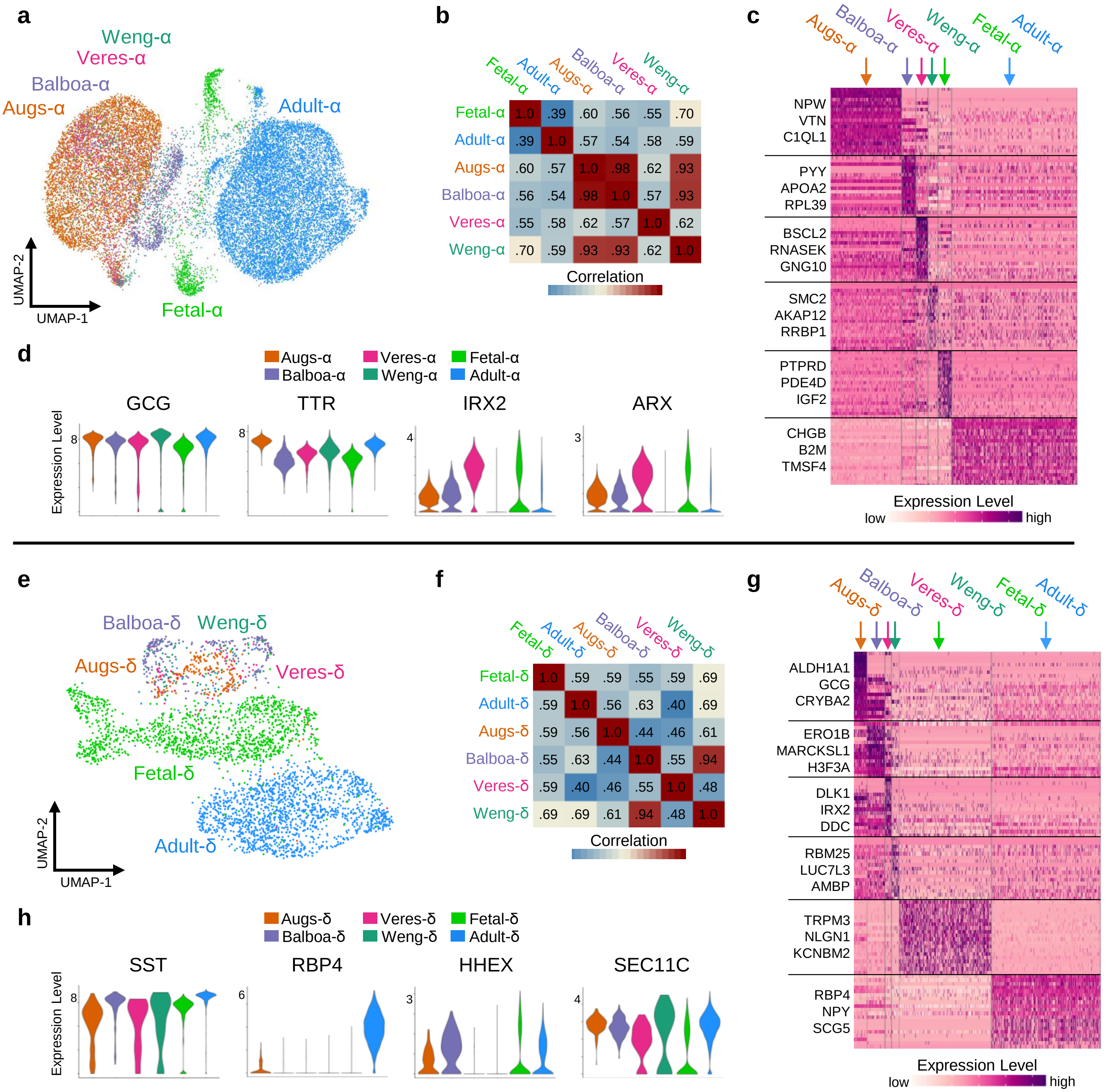


**Fig. S1 | SC-α and SC-δ cells derived from different protocols have distinct transcriptional differences compared to their adult counterparts, related to Fig 2.** (a) UMAP of adult α-cells, fetal α-cells, and SC-α cells clustered and split by their derivation protocol. (b) Heatmap of Pearson correlation coefficient for 1000 most variable expressed genes. (c) Heatmap of scaled RNA expression indicating top 20 most differentially expressed genes for α-cells derived by unique protocols, fetal α-cells, and adult α-cells. (d) Violin plots indicating expression level of α-cell identity markers. (e) UMAP of adult δ-cells, fetal δ-cells, and SC-δ cells clustered and split by their derivation protocol. (f) Heatmap of Pearson correlation coefficient for 1000 most variable expressed genes. (g) Heatmap of scaled RNA expression indicating top 20 most differentially expressed genes for δ-cells derived by unique protocols, fetal δ-cells, and adult δ-cells. (h) Violin plots indicating expression level of δ-cell identity markers.

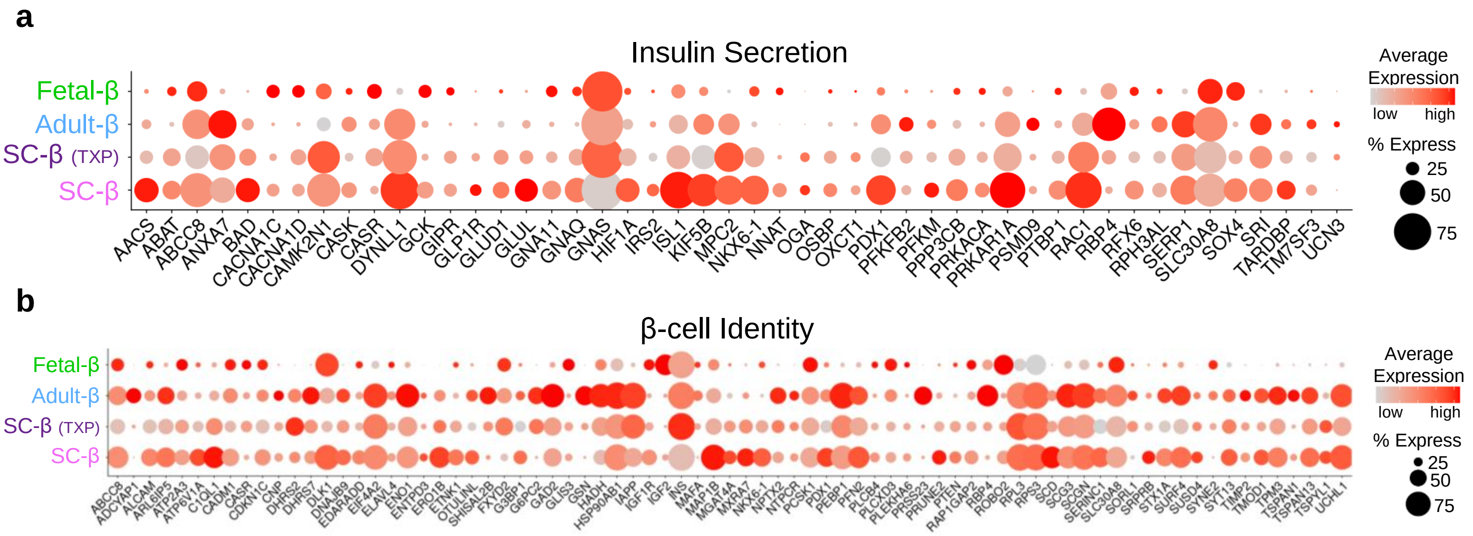


**Fig. S2 | Fetal β-cells lack expression of important β-cell machinery, related to Fig 3.** (a) Dot plot indicating proportion of cells expressing and average expression of genes involved in insulin secretion. (b) Dot plot indicating proportion of cells expressing and average expression of genes involved in β-cell Identity.

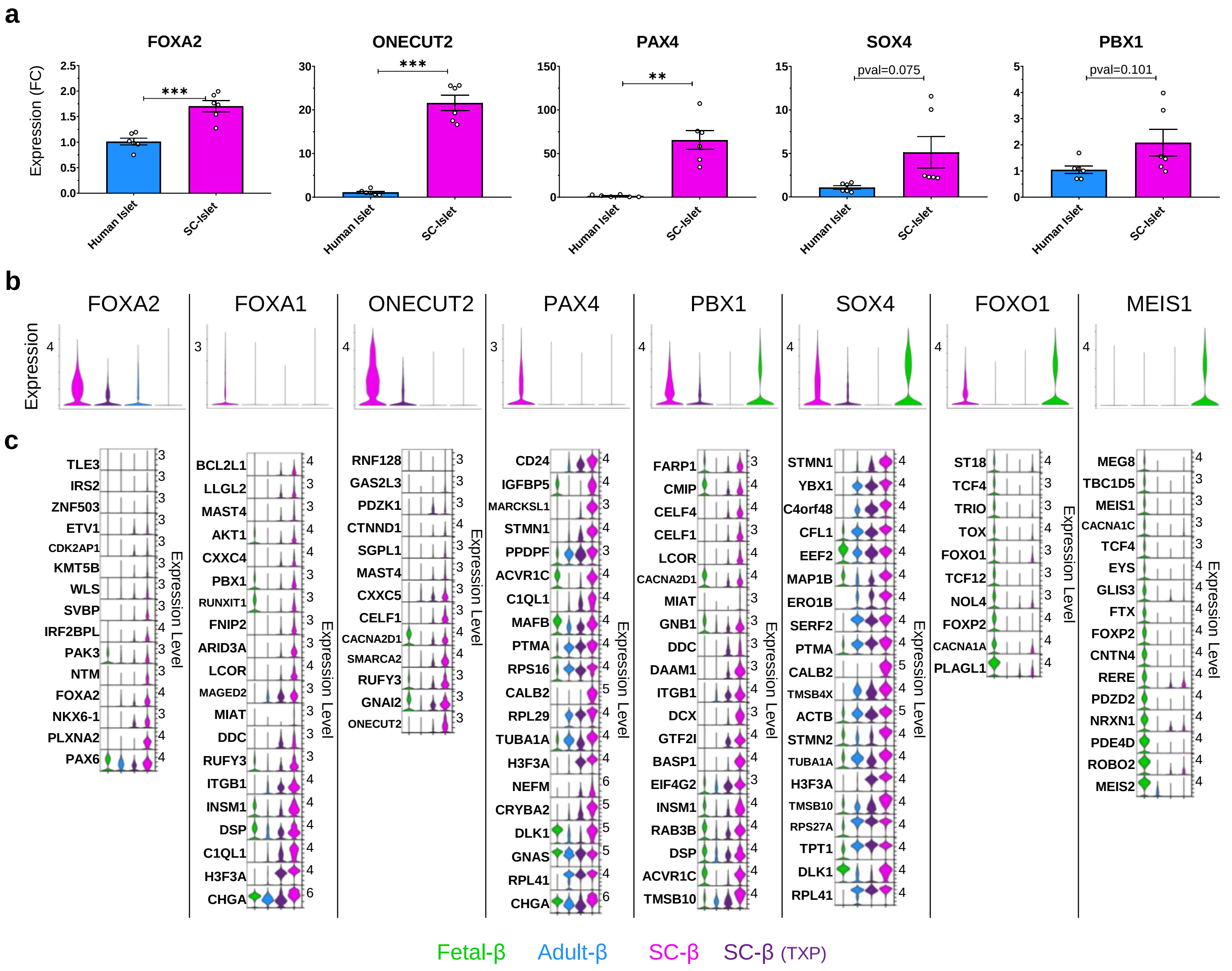


**Fig. S3 | Validation of enriched regulons in SC and fetal β-cells through expression of transcription factor target genes, related to Fig 4.** (a) RT-qPCR of SC-islets (n=6) at s6d7 and human cadaveric islets (n=6) indicating fold change (FC) expression relative to TBP. (b) Violin plots indicating expression level of transcription factors that have enriched regulon activity. (c) Violin plots indicating expression level of highly expressed transcription factors targets associated with indicated regulon. All data are represented as the mean, and all error bars represent the s.e.m. Individual data points are shown for all bar graphs. ns, not significant; *P< 0.05, **P< 0.01, ***P< 0.001.

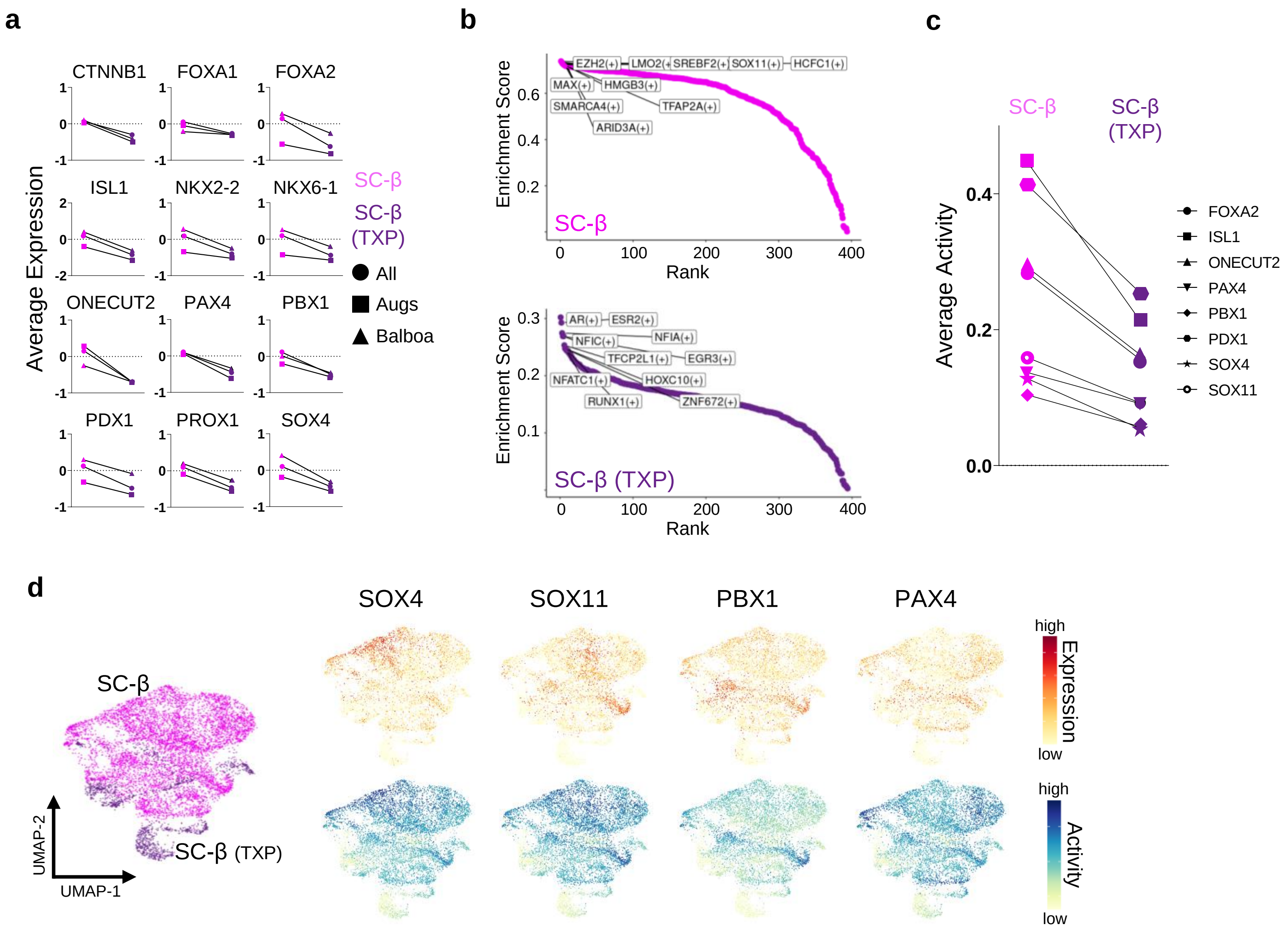


**Fig. S4 | Expression and activity of progenitor transcription factors decreases in SC-β cells after transplantation, related to Fig 4.** (a) Line graphs indicating decrease in average expression of known transcription factors associated with β-cell development after transplantation of SC-β cells. (b) Chart indicating highest predicted active regulons in SC-β versus transplanted SC-β cells. (c) Line graph indicating decrease in average activity of known β-cell progenitor transcription factors after transplantation of SC-β cells. (d) UMAP of SC-β and SC-β TXP cells clustered by RNA expression and feature plots indicating expression and activity of β-progenitor transcription factors.

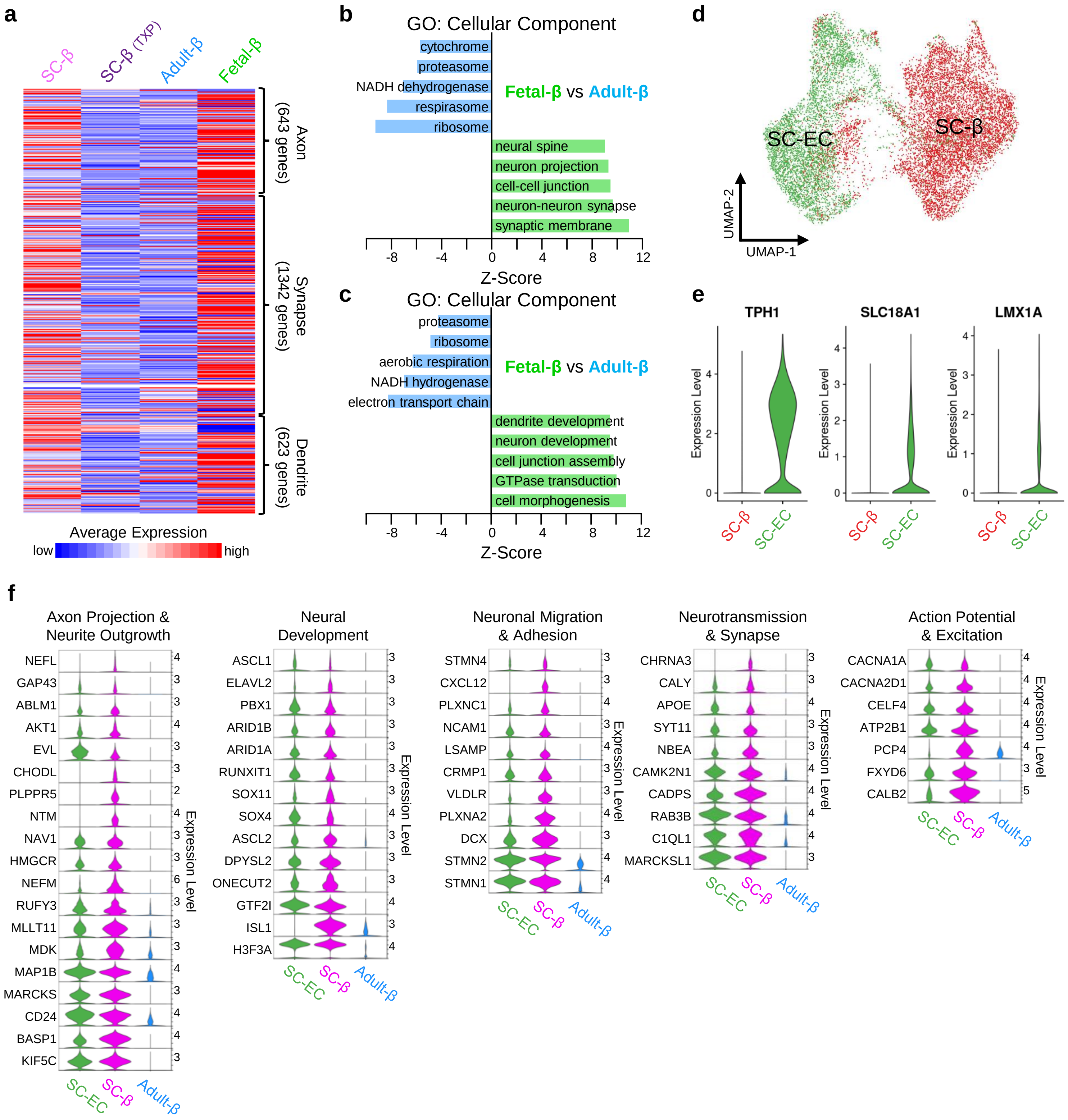


**Fig. S5 | Transcripts involved in synaptic function are enriched in SC-β, fetal β-cells, and SC-EC cells, related to Fig 5.** (a) Heatmap indicating average expression of genes associated with axon, synapse, and dendrite formation between SC, SC-TXP, adult, and fetal β-cells. (b) Bar chart indicating gene ontology: cellular component enrichment scoring of differentially expressed genes between fetal-β and adult-β cells. (c) Bar chart indicating gene ontology: biological process enrichment scoring of differentially expressed genes between fetal-β and adult-β cells. (d) Integrated UMAP of all SC/SC-TXP β and EC cells. (e) Violin plots indicating expression level of genes associated with serotonin machinery in SC/SC-TXP β and EC cells. (f) Panel of violin plots indicating expression level of genes associated with various neuronal traits between SC-β, SC-EC, and adult-β cells.

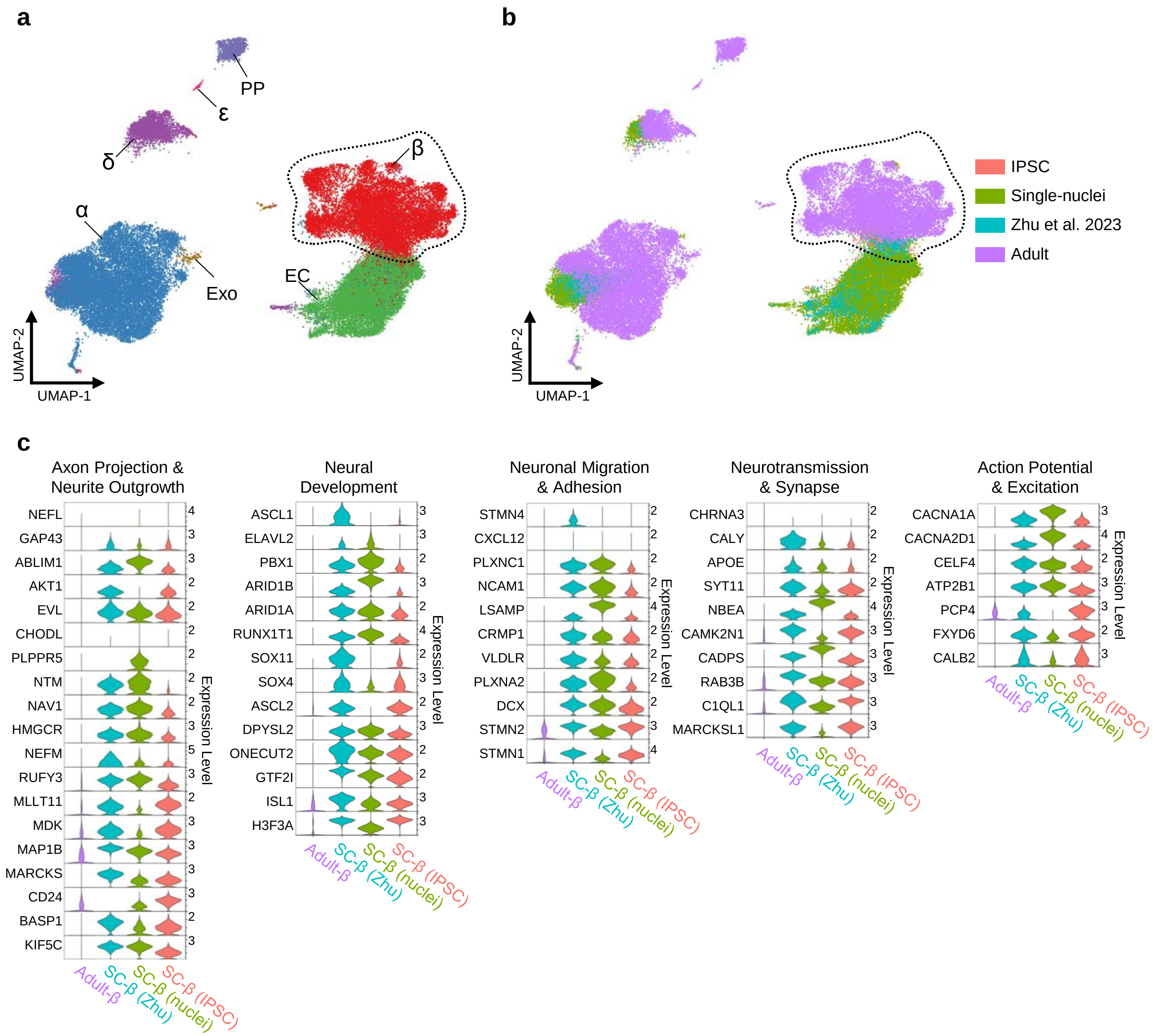


**Fig. S6 | Validation of neural gene program using additional datasets, related to Fig 5.** (a) Integrated endocrine islet UMAP of cells from SC-islets derived from an IPSC cell-line (Augsornworawat et al. 2020), SC-islets derived from the H1 hESC cell-line (Zhu et al. 2023), SC-islets derived from the Hues8 hESC cell-line and sequenced using single-nuclei RNAseq method (Augsornworawat et al. 2023), and human adult islet cells. 7 cell types were identified. Enterochromaffin-like (EC), Exocrine (Exo). (b) Integrated endocrine islet UMAP grouped by their source. (c) Panel of violin plots indicating expression level of genes associated with various neuronal traits between adult β-cells and populations of SC-β derived with unique cell-lines and sequencing platforms.

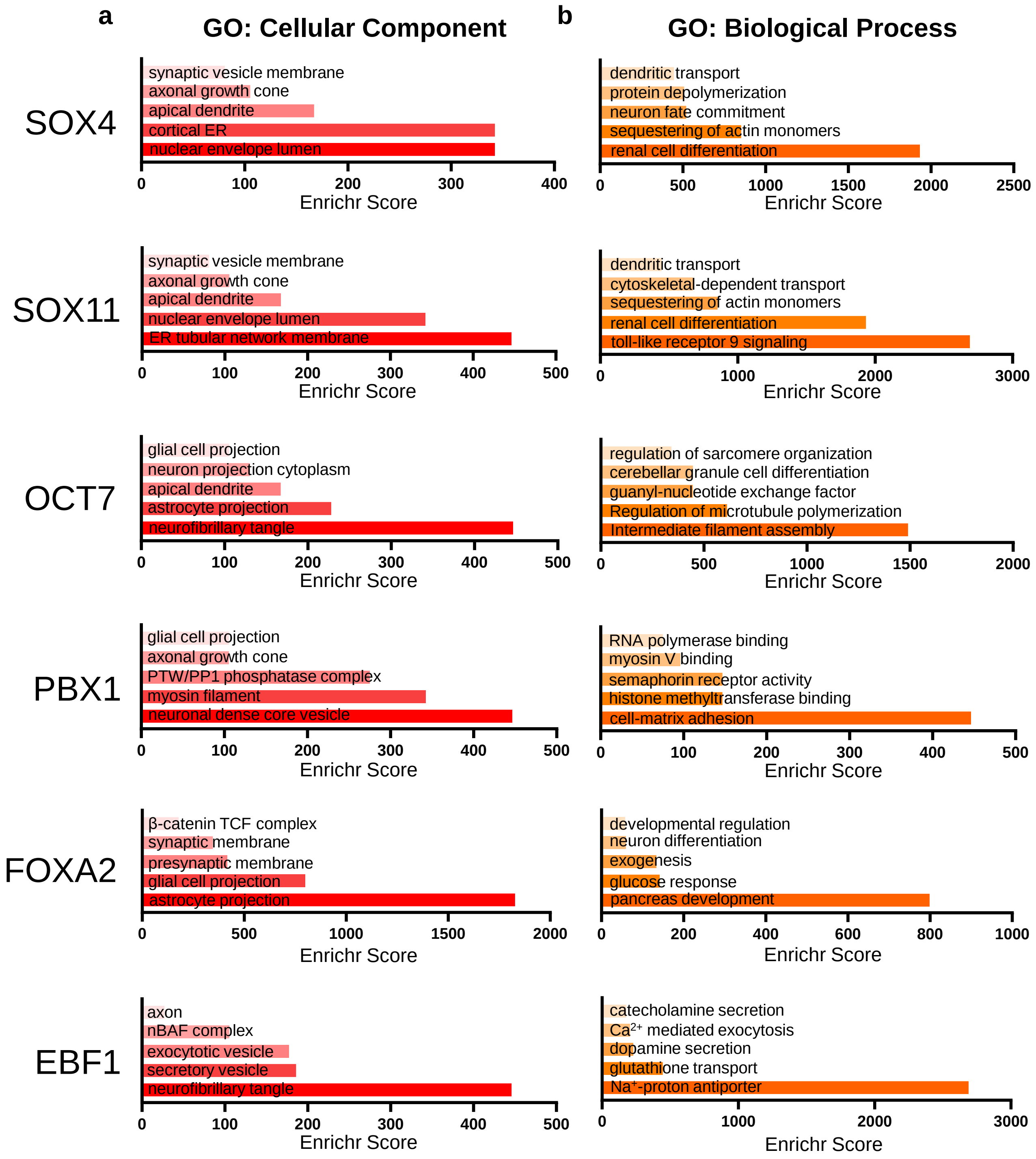


**Fig. S7 | EnrichR identifies neuronal morphological and biological gene sets enriched in gene targets of implicated transcription factors, related to Fig 6.** (a) High Enrichr scoring GO: Cellular Component terms associated with the top-50 expressed gene targets of indicated regulon. (b) High Enrichr scoring GO: Biological Process terms associated with the top-50 expressed gene targets of indicated regulon.
